# A Network Medicine Approach to Drug Repurposing for Chronic Pancreatitis

**DOI:** 10.1101/2020.10.30.360263

**Authors:** Megan Golden, Jabe Wilson

## Abstract

Despite decades of clinical investigations, there is currently no effective treatment for patients diagnosed with Chronic Pancreatitis (CP). Computational drug repurposing holds promise to rapidly identify therapeutics which may prove efficacious against the disease. Using a literature-derived knowledge graph, we train multiple machine learning models using embeddings based on i) the network topology of regulation bipartite networks, ii) protein primary structures and iii) molecule substructures. Using these models, we predict approved drugs that down-regulate the disease, and assess their proposed respective drug targets and mechanism of actions. We analyse the highest predicted drugs and find a diverse range of regulatory mechanisms including inhibition of fibrosis, inflammation, immmune response, oxidative stress and calcium homeostasis. Notably, we identify resiniferatoxin, a potent analogue of capsaicin, as a promising repurposable candidate due to its antiinflammatory properties, nociceptive pain suppression, and regulation of calcium homeostatis (through potentiation of mutant cystic fibrosis transmembrane conductance regulator (CFTR)). Resiniferatoxin may also regulate intracellular acinar Ca2+ via agonism of transient receptor potential vanilloid subfamily member 6 (TRPV6). We believe the potential of this repurposable drug warrants further *in silico* and *in vitro* testing, particularly the affect of the TRPV6 agonism on disease pathogenesis.

## 1 Introduction

TO date, there is no approved and satisfactory means of treatment for Chronic Pancreatitis (CP). Despite advances in mechanistic understanding of the disease, no drugs have been developed that satisfactorily prevent either abdominal pain or prevent disease progression. Repurposed drugs hold great promise as potential treatment due to their established safety profiles and established manufacturing routes; minimising time to use these drugs at the point-of-care.

Systems pharmacology and network medicine (NM) approaches to drug discovery and drug repurposing have proved to be efficient for the identification of potential drug candidates [1], [2], [3]. NM treats biological networks as heterogeneous information systems; correlating network topology and node properties with biological processes, functions, pathways and interactions. From a systems biology point of view, a disease can be seen as a selection of genes within a network, whose dysregulation culminates in changes in biological processes, pathways and ultimately phenotype. Similarly, drugs can be modelled by their drug targets and the propagatory effect their perturbance has upon the network. The aim of NM is to identify drugs and diseases in which the network perturbance of the disease state is reversed by the perturbance of the drug.

The interactome (the totality of interactions within a cell), is largely unknown and fascinatingly complex. Drug target interactions constitute much less than one percent of small-molecules reported to bind to a protein. Drug regulatory interactions with disease follow a similar distribution. Our sparsity in known information holds promise that there are novel indications for current drugs; through the reapplication of drugs to new diseases, through both known and novel targets.

In recent years, graph-based machine learning (GML) methods have been applied to systemically ‘fill in’ these unreported interactions through the process of link prediction. Whilst many GML methods exists, the most intuitive is graph neural networks. In contrast to conventional neural networks which use arbitrary model architectures, GML models explicitly replicate the network relating to their prediction task. For example, using the aforementioned disease-gene network, and tasked to predict novel gene regulators of disease, genes which are functionally or physically similar are more highly connected within the biological network, and by extension are more connected in the model architecture. Such genes will have greater influence on each other during the training process of the model. One of the applications of GML is the creation of node embeddings: transforming continuous high dimensional information of node neighbourhood to low-dimensional dense vectors, to be used in downstream machine learning tasks. Popular approaches include generating embeddings by sampling a network via random walks, treating the sample as corpus of words, and applying natural language processing (NLP) techniques. Similar NLP methods have been applied to amino acid sequences of proteins [4], and molecular substructures of compounds [5] to generate embeddings representing primary structure and molecular substructure of proteins and compounds respectively.

## 2 Results

### 2.1 Knowledge Graph

To model CP from a systems biology point of view, we need to create a network capable of capturing the biochemical and regulatory interactions involved in disease progression. We created a knowledge graph predominantly based on Pathway Studio, a biomedical database derived from relationships extracted from over 30 million academic manuscripts (see Methods). Each edge in the graph represents at least one occurrence in scientific literature that has stated a relationship between biological entities (that ibuprofen down-regulates acute pancreatitis). In total, the graph possessed 1.36 million nodes and 8.52 million edges, weighted according to their frequency in literature (see Fig 1). From this we generated a tripartite subgraph of diseases, drugs and genes. We included three edges types: inhibition and activation (via any direct or indirect mechanism) and regulation; a conflation of up-and down-regulation and regulation of unknown direction (see Figure 2).

**Fig. 1.**
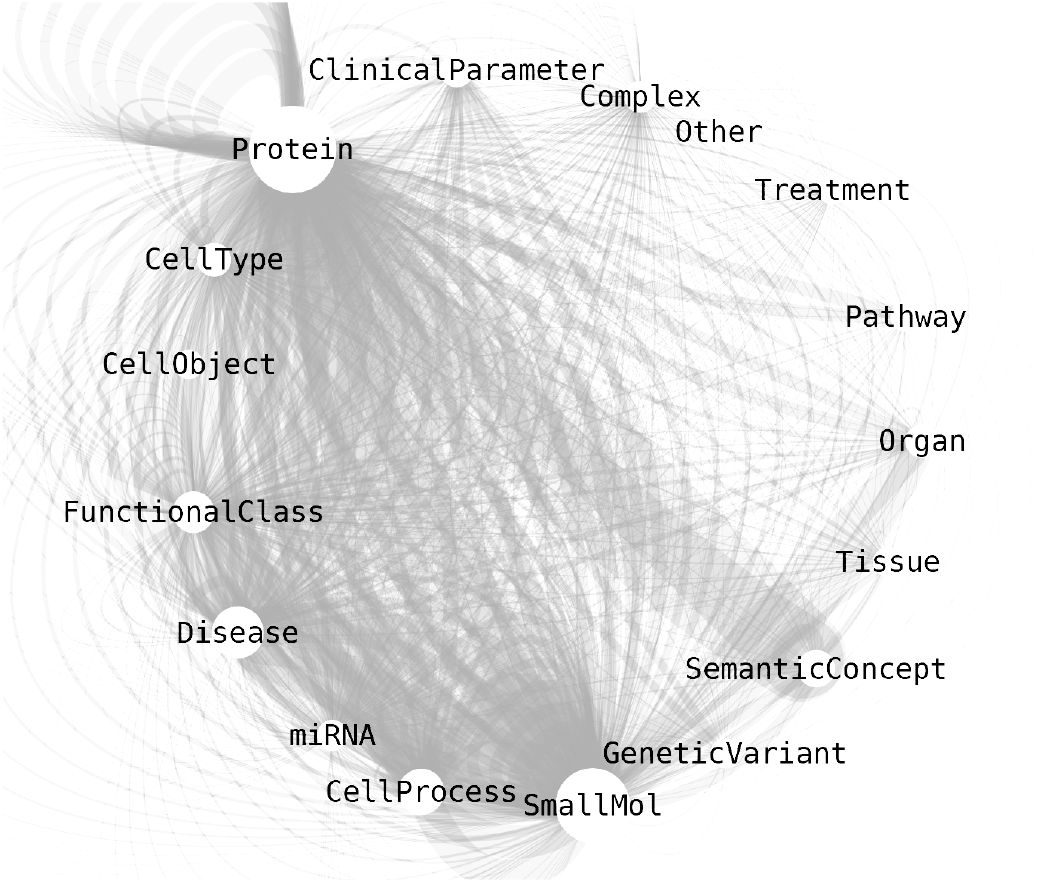
Cytoscape visualisation of the knowledge graph. Size of nodes represent number of occurrences in the graph

**Fig. 2.**
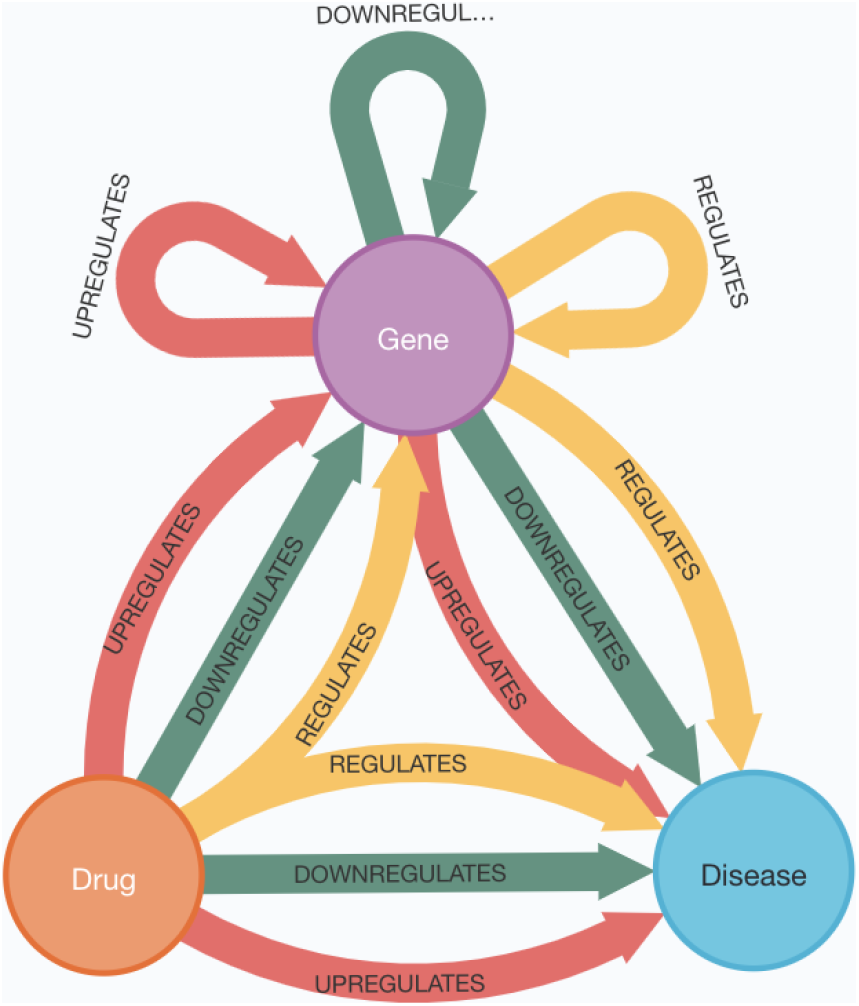
Graph database schema.

### 2.2 Predictions

#### 2.2.1 Regulators

To ascertain the therapeutic efficacy of drugs for treatment of CP, we developed an embedding-based link prediction model capable of predicting the existence of a relationship (otherwise known as an edge) between two nodes. We trained link prediction models to predict drugs that inhibit any disease node (see Table 1). As this edge is based on occurrence of biological relationships in literature, a predicted link with high probability for the edge *ibuprofen-inhibits-chronic pancreatitis* indicates that if a research group were to research the inhibitory regulation of the drug upon the disease, there is high likelihood the regulation was sufficient to state this regulatory relationship in an academic paper. The top 25 predictions for CP can be seen in Table 3. For each prediction, we cross-referenced and highlighted known down-regulators reported in literature, and appended known drug targets of each respective drug.

**TABLE 1.** Models used in analysis. *Edge* column describes the link being predicted, from which one of the three embeddings was generated. The *Source* and *Target* columns describe the type of embeddings used for the additional respective source and target features. Regulates abbreviated as *reg*.

| Name | Edge | Source | Target |
| --- | --- | --- | --- |
| CdD | Drug-down-reg-disease | Mol2Vec | Gene-reg-disease |
| GdD | Gene-down-reg-disease | ProtVec | Drug-reg-disease |
| GuD | Gene-up-reg-disease | ProtVec | Drug-reg-disease |

Our model used node embeddings to create compressed discrete 100-dimensional representations of the neighbourhood of each node. After various evaluations, we chose to employ the state-of-the-art embedding method, GraRep [6], due to the method’s ability to capture both local and global neighbourhood information. In essence, GraRep calculates the singular value decomposition for direct neighbours of a node, and indirect neighbours up to a certain threshold distance. For a full explanation, please see the original paper. For an illustrative explanation of the link prediction model, please see the Methods section.

Alongside information of node neighbourhood, the model also utilised physicochemical structural information of proteins and molecules. We used Mol2Vec [5], an unsupervised machine learning approach to learn vector representations of molecular substructures, to capture structural information of drugs. ProtVec [4] was used to represent the primary amino acid structure of proteins. For disease nodes, additional graph embeddings were generated based on a disease-gene regulatory and disease-drug regulatory networks for the drug-down-regulates-disease (CdD), gene-down-regulates-disease (GdD) and gene-up-regulates-disease (GuD) models. A high-level overview of model performance can be seen in Table 2).

**TABLE 2.** Model performance over 5 random folds. Standard deviation shown. *Name* abbreviation refers to models in Table 1.

| Name | AUC | F1 | Accuracy |
| --- | --- | --- | --- |
| CdD | 0.936±0.002 | 0.888±0.003 | 0.888±0.003 |
| GdD | 0.954±0.002 | 0.868±0.004 | 0.870±0.004 |
| GuD | 0.927±0.002 | 0.879±0.001 | 0.876±0.003 |

## 3 Discussion

### 3.1 Notable Drug Candidates

Chronic Pancreatitis is a fibro-inflammatory disease, primarily caused by pancreatic duct obstruction initiating intracellular activation of pancreatic proenzymes and autodigestion of the pancreas. We used a machine learning model to predict the likelihood of drugs to down-regulate CP. The top 25 predictions (see Table 3) demonstrate the heterogeneity of the disease, its etiologies, and respective mechanistic pathways. A brief explanation of notable candidates and their supposed mechanism of action is discussed below.

**TABLE 3.** Top 20 predicted inhibitors of CP. PubMed IDs are provided for known relationships. † indicates drug is or was in clinical trial for CP.

| Drug | Prob | PMID of edge | Drug Class | Drug Targets |
| --- | --- | --- | --- | --- |
| Capsaicin | 0.9896 | 21859833, 22350613 | Analgesic | TRPV1, TRPV6 |
| Estradiol | 0.9880 | 12657966 | Endogenous hormone | ESR1, ESR2 |
| Pioglitazone† | 0.9844 | 17510194 | Antihyperglycemic | PPARγ |
| Dimethyl fumarate | 0.9824 | 25198679, 26972398, 27499754 | Immunosuppressant | KEAP1 |
| N-acetylcysteine | 0.9820 | 7922442 | Mucolytic | GSS, SLC7A11 |
| Ibuprofen | 0.9820 | 22918205 | NSAID | PTGS1, PTGS2 |
| Pentoxifylline | 0.9772 | - | Hemorrhologic agent | ADORA1, ADORA2A |
| Nicotinamide | 0.9768 | - | Antioxidant | - |
| Ellagic acid | 0.9764 | 18651219 | Antioxidant | - |
| Fingolimod | 0.9764 | 15782091 | Immunosuppressant | S1PR1,S1PR3,S1PR4,S1PR5 |
| Minocycline | 0.9764 | 21963786 | Antibacterial | - |
| Rofecoxib | 0.9748 | 21372163, 28551708 | NSAID | PTGS2 |
| Celecoxib | 0.9744 | - | NSAID | PTGS2 |
| Quercetin | 0.9744 | - | Antioxidant | - |
| Caffeine | 0.9724 | - | Psychoanaleptic | ADORA1,ADORA2A,ADORA2B,ADORA3 |
| Dopamine | 0.9724 | - | Cardiac stimulant | DRD1, DRD2, DRD3, DRD4, DRD5, SLC6A3, DBH |
| Naproxen | 0.9716 | 19823098 | NSAID | PTGS1, PTGS2 |
| Pravastatin | 0.9716 | 21383674, 26773928, 28551708 | Lipid modifying agent | HMGCR |
| Tacrolimus | 0.9712 | 15782091 | Immunosuppressant | FKBP1A |
| Ascorbic acid | 0.9704 | 27306367 | Antioxidant | - |
| Indomethacin† | 0.9700 | - | NSAID | PTGS1,PTGS2,PLA2G2A |
| Metformin | 0.9700 | - | Antihyperglycemic | PRKAB1, ETFDH |
| Fenofibrate | 0.9696 | - | Lipid modifying agent | PPARα |
| Melatonin | 0.9696 | - | Psycholeptic | MTNR1A, MTNR1B |
| Phenylbutyric acid | 0.9688 | - | Chemical chaperone | SP-A2, A1ATD, CFTR |

The most common symptom of chronic pancreatitis is repeated episodes of severe abdominal pain. Both analgesics and non-steroidal anti-inflammatory drugs (NSAIDs) have been historically administered to alleviate such pain. Many NSAIDs such as ibuprofen, naproxen, indomethacin and celecoxib were highly predicted to modulate CP. There are however conflicting reports that NSAIDs may induce acute pancreatitis (AP). One study showed an increased risk of AP for all NSAIDs: lowest for the naproxen-treated group. The study also highlighted those treated with indomethacin (the only NSAID currently in clinical trial for CP) showed lower risk of post-endoscopic cholangiopancreatography (ERCP) AP and less gastrointestinal bleeding [7]. CP is characterised by destruction of the pancreatic parenchyma, first inducing a local inflammatory reaction, leading to overwhelming systemic production of inflammatory mediators and early organ failure [8]. It is has been suggested that the antiinflammatory properties of NSAIDs could reduce systemic complications and ensuing tissue damage. Whilst the situation remains unclear, a systematic review of 36 clinical studies clinical evidence showed that NSAIDs are effective in suppressing proinflammatory cytokines, ameliorating systematic complications and reducing mortality (alongside their predominant purpose of reducing pain) [9]. Such a result provides therapeutic promise for NSAIDs for treatment of CP.

Capsaicin was the highest predicted drug to treat CP. Literature concerning the affect of the botanical upon CP is conflicting. Researchers have reported that capsaicin significantly reduced the severity of chronic pancreatitis, as determined by evaluating the loss of acini, inflammatory cell infiltration and stromal fibrosis [10]. Known to have antiinflammatory properties, this clinical improvement was attributed to a significant decrease in inflammatory cells including neutrophils and macrophages in the pancreas. Capsaicin has also been shown to reduce tissue damage in experimental acute pancreatitis by releasing endogenous calcitonin gene-related peptide, improving pancreatic microcirculation and reducing inflammation [11]. Capsaicin is a selective agonist of TRPV1 and TRPV6. TRPV1 antagonism has showed decreased visceral pain behaviour in mice models [12], indicating visceral nerve sensitization via TRPV1 antagonists may be a therapeutic option for CP. Capsaicin has been shown to activate and sensitise pancreatic nociceptors and increase neuropathic pain [13]. Conversely, excessive TRPV1 agonism is known to lead to cytotoxic calcium overload and cell death of TRPV1-positive neurons [14]. Resiniferatoxin (RTX) (a naturally-occurring ultrapotent capsacain analogue and TRPV1 agonist) is currently in multiple clinical trials for urinary bladder hyper-reflexia and chronic pain conditions such as osteoarthritis. Administration of RTX reduced severity of AP [15], defunctionalising the nociceptive nerve fibres C and Adelta and modulating the TRPV1-dependent mechanism of (post-ECRP) pancreatitis [16]. This may explain the lack of agreement, as agonism of TRPV1 sensitises and exacerbates pain, whilst sufficiently potent agonism defunctionalises the nerve fibres, attenuating the pain.

Due to the ubiquity in which TRPV1 is expressed throughout the central nervous system, researchers have suggested local delivery of agonists to TRPV1-positive nociceptors, to alleviate pain without systemic side effects [17]. Patent literature describes the application of RTX to a visceral organ such as the stomach or jejunum (organs that share a spinal or vagal nerve with the pancreas) may modulate pain in the pancreas, alongside inhibiting inflammation [18]. This may prove a viable delivery route of TRPV1 agonists.

Loss of function mutations in CFTR are widely regarded as the predominant pathomechanism of CP via dysregulation of ductal Ca2+. Ivacaftor, a CFTR potentiator, has seen most clinical success in treatment of cystic fibrosis. Capsaicin is known to potentiate both wild-type and mutant (G551D-CFTR, ΔF508-CFTR, and 8SA) CFTR channels [19]. A recent study demonstrated functionally defective variants in TRPV6 were associated with early onset CP in a small subpopulation of patients, hypothesised to be due to dysregulation of Ca2+ homeostasis [20]. The research group demonstrated loss of function of TRPV6 increased intracellular Ca2+, which was correlated with early onset CP. Both capsaicin and RTX are potent TRPV6 agonists and thus may have an affect on restoration of intracellular calcium levels. RTX has certainly cemented itself as a therapeutic candidate due to its potent and selective neuropathic pain suppression, inhibition of inflammation, and regulation of calcium homeostasis via CFTR potentiation. Its agonism of TRPV6 and affect on ductal Ca2+ needs to be assessed.

Oxidative, electrophilic, ER, and inflammation stress are widely regarded to contribute to the pathogenesis of CP [13]. Micronutrient therapy has been suggested to inhibit various etiological stresses [21]. Indeed, many antioxidants such as N-acetlycysteine, nicotinamide, ellagic acid and quercetin were highly predicted to regulate CP. N-acetlycysteine (NAC), a reduced glutathione provider and the fifth highest predicted therapeutic, is known to inhibit oxidative stress [22] and endoplasmic reticulum (ER) stress [23], [24]. The reduction of oxidative-stress-induced cell injury has highlighted NAC as a potential treatment of CP [25]. NAC acts as a direct scavenger of reactive oxygen intermediates, preventing production of oxygen free radicals (OFRs) [25]. One of the predominant etiological factors of disease progression is a build up of cytosolic Ca2+. OFRs are known to disturb calcium homeostatis. Administration of NAC has been shown to inhibit the increase of cytosolic Ca2+ in acinar cells, ultimately reducing the accumulation of proenzymes and slowing disease progression [25]. Alongside NACs ability to restore calcium homeostasis, the mucolytic may exert an antiinflammatory effect through the inhibition of tumor necrosis factor-*α* (TNF-*α*), decreasing the plasma level of interleukin (IL)-6 and blocking nuclear factor-kappaB (NF-*κ*B) activation [26].

Other highly predicted micronutrients included nicotinamide adenine dinucleotide (NAD^+^). Increase of NAD^+^, an oxidative agent involved in myriad cellular processes, was shown to inhibit TLR4: reducing the inflammation and cell death [27]. Ellagic acid, also highly predicted, has been shown to attentuate pancreatic inflammation and fibrosis via the inhibition of reactive oxygen species production in profibrogenic pancreatic stellate cells [28]. High dose vitamin C (ascorbic acid) demonstrated therapeutic efficacy against AP, namely due to the micronutrients anti-oxidizing ability [29].

Pentoxifylline (PTX) is a non-selective phosphodiesterase inhibitor with antiinflammatory, antifibrotic and antioxidant properties. TNF-*α* is a key component in CP progression. PTX non-selectively antagonises the inflammatory cytokine. As such, PTX has been applied to various other TNF-*α*-modulated diseases including AP, and has shown to be well-tolerated and efficacious [30]. PTX inhibits platelet derived growth factor and other cytokines essential to fibrotic processes. Due to this antifibrotic effect, PTX has also been repurposed to an assortment of conditions such as oral submucous fibrosis [31] and radiation-induced fibrosis [32]. It has been suggested that PTX could slow the course of pancreatitic fibrosis [33].

An intriguing prediction was lipid modifying agents pravastin and fenofibrate. Historically, statins have largley been associated to pancreatitis by their supposed induction of the disease. Recently the prevalence of pancreatitis secondary to statins has been questioned. Only 12 reports of statin-induced pancreatitis have been reported [34]. More recent studies have demonstrated that statins may actually have a protective role for CP, attenuating inflammation and fibrosis, most likely due to their antioxidative properties [35].

### 3.2 Methodology

As with all *in silico* methodologies, there are many limitations to this workflow. An important note is that the above diseases have vastly differing degrees of connectivity in the knowledge graph. Whilst a well-researched disease such as pancreatic cancer have 100s of connected down-regulators in the knowledge graph, the number of reported inhibitors for chronic pancreatitis is an order of magnitude fewer. Such differences in degree indicate different prior probabilities of treatment. In other words, nodes in a graph with higher connectivity are statistically more likely to be connected. This prior probability is reflected in the probability distributions. Pancreatic cancer will have a considerably higher average probability compared to CP. The same applies to any type of node in a link prediction problem. Known genes (notably oncogenes and tumor suppression genes) and popularly used drugs (such as NSAIDs and immunosupressants) will have significantly higher prior probabilities, appearing higher in the prediction lists, even if the local topological evidence between a source and target node is less.

Link prediction on knowledge graphs also signifies that the area under the receiver operating characteristic curve is not 0.5 for random guesses of connecting source and target nodes: it is much higher. Researchers have shown prior probability of treatment (the likelihood two nodes are connected simply by their node degrees) can be achieved by randomly permuting bipartite graphs multiple times (swapping edges but preserving node degree) and noting the average probability of an edge as a function of source and target degree. Ensuring an edge prediction is larger than the prior probability of connection, guarantees local network topology suggests the presence of an unreported edge, not simply the global degree distribution. The prior probability dominates predictions on most networks. One recent attempt to overcome this uses a Bayesian approach during the training process of the graph embeddings; modelling the prior probability as the prior in the Bayes formula [36]. By explicitly modelling the prior, it ensures it is not captured in the graph embedding.

Machine learning methods are often cursed by their lack of explainability. Specifically, the optimized parameters of supervised models are hard to interpret. Whilst embeddings methods have proved powerful strategies to capture continuous data into discrete and dense vector representations, they obfuscate explainability even further. Embeddings are used as the features of a model. As each dimension is the output of another vectorization model, they are completely uninterpretable.

Biomedical knowledge graphs are largely incomplete. On average, well under one percent of possible relationships are reported for edges such as regulation, binding and expression. This presents a problem for link prediction models; the true distribution of positive and negative classes is unknown, and it is impossible to differentiate a true (but unreported) positive from a false positive. It is preferable to train your model on a class balance that matches your real world distribution. In this application, this is not possible. Moreover, when your real world distribution is known, optimizing your classification threshold becomes much simpler task. We used maximal f1 score to determine optimal threshold. This often meant an unrealistic increase in the number of positive predicted samples. For 3302 drugs and 19387 diseases, there are 163590 known inhibitors (0.25 percent). The average positive class for a random subset of predicted regulators above the optimal threshold of 0.634 was 28 percent, a 112-fold increase in regulators. This is obviously a gross overestimation in the number of unreported regulators and a major limitation in our approach. We encourage predictions to be analysed only on the context of the relative list of regulators for that disease. An alternative approach taken to decide classification threshold would be based largely on the known class distribution [2].

The knowledge graph provides an incredible amount of biological information, and we only used an incredibly small amount of this information. Each edge in the graph was weighted according the absolute occurrence that this relationship appears in scientific literature. It it logical to believe the number of times a relationship is reported is correlated with its strength or confidence. This does, however, introduce a new level of knowledge bias. Genes such as p53 have received tremendous attention. Does this signify said gene’s importance over it’s less-researched genetic counterparts? In previous unpublished analyses, we attempted to utilise this weighting through numerous normalisation strategies such as frequency of node, edge and path and log-scaled reference counts. We found that such strategies only generalised solutions; wherein well-researched areas performed better, whilst less-researched performed worse. An interesting approach would be to normalise nodes according to the weighted edges that connect to it (weighted by reference count). In this case, for a well-researched node with 10 edges and an individual edge weight of 50, each edge would be equivalent to a node with 10 edges and an edge weight of 5. This may also solve the prior probability issue stated earlier, as for nodes with fewer edges, each edge will be given higher importance, partially mitigating the dominating effect of nodes with high degrees.

## Conclusion

There are currently no satisfactory means of treatment of CP. Using a network pharmacology approach, we highlighted the potential therapeutics, many of which have been previously investigated regarding their affect on acute pancreatitis or chronic pancreatitis. Of the top contenders, we believe resiniferatoxin warrants the most attention due to its known antiinflammatory properties, known visceral pain suppression, and potential regulation of Ca2+ homeostatis between acinar cells and ducts. We suggest the pharmacological viability and therapeutic efficacy of agonism of TRP channels should be further investigated *in vitro*, and the required pharmacokinetics assessed.

## 4 Methods

### 4.1 Drug Regulators

#### 4.1.1 Knowledge Graph

We created a knowledge graph predominantly based on Pathway Studio, a literature-derived database that uses natural language processing techniques to leverage biological relationships from over 30 million literary sources. Pathway Studio also contains the relevant subset of Reaxys Medicinal Chemistry, a database of small-molecule protein bioactivities, pertaining the species homo sapiens, Mus muccus, and Rattus rattus, and rattus norvegicus. We appended this core graph with gene ontologies [37], drug side effects from SIDER [38] and drug-target information from DrugCentral [39]. We also created similarity links such as protein-protein similarity (Local Smith Waterman of over 0.5), and molecule substructure similarity (Tanimoto similarity of Morgan Fingerprint of over 0.5). After refactoring and harmonization, the graph possessed 1.36 million nodes and 8.52 million edges and over 200 edge types, weighted according to their occurrences in literature.

For each molecule with an InChI code or key within the graph, we generated a Mol2Vec embedding using the pretrained embeddings [5] were trained on 20M compounds form the ZINC database [40]. As all other embeddings had a dimension of 100, and as the model requires embeddings of the same size, we used the scikit-learn version of principal component analysis to reduce the embedding down to the required size. Summation of the explained variance showed only 1.3 percent of information was lost. Similarly, we generated embeddings for proteins based on trimers of their amino acid sequence using the pretrained model of ProtVec [4], trained on 551,754 proteins from Swiss-Prot. Because trimers can start at the first, second or third amino acid in a protein sequence, three embeddings were generated per protein. As per the methodology of the original paper, we took the element-wise average of these.

For this analysis, we created a tripartite subgraph of diseases, genes and drugs, connected via multiple regulatory edges. Said edges included up- and down-regulation via any direct or indirect mechanism. For example, the edge *drug-inhibits-gene* conflates expression, indirect regulation, direct binding (agonism and antagonism), and promoter binding. We also provided a conflated edge of both up- and down-regulation.

#### 4.1.2 Regulation Prediction

To determine if a drug down-regulated a disease or gene, we developed an embedding-based link prediction model based on multiple disease regulatory bipartite networks and additional physicochemical and structural information of the source and target nodes (see Fig 3). The embeddings were used by a random forest classifier (scikit-learn implementation), optimized via hyper-parameter bayesian optimization. We assessed multiple node embedding strategies in this project. For embedding choice in the link prediction model, we assessed GraRep, nodevec, LINE and SVD model. All models used the BioNev [41] implementation, except node2vec, for which we used the C++ version from SNAP [42]. Random search was employed to determine models with the highest AUC score. We also investigated different mathematical functions to create an edge embedding from two node embeddings (concatenation, element-wise average, hadamard, L1 and and L2 loss). Because our model used two different types of embeddings for each edge (for example: i) graph embedding for protein and molecule, and ii) ProtVec and Mol2vec respectively), employing the hadamard edge function signifies, the hadamard was calculated both for graph embeddings, and the for Mol2vec and ProtVec separately before concatenating. We also investigated stacking models to create soft-voting bagging classifiers. Because a subset of edges must be removed and used to train the model, we postulated that training multiple models on different subsets, and combining their predictions via a weighted average according to the performance, would increase predictive power of the stacked model. Results showed, however, that increase was extremely marginal (under 0.5 percent increase in AUC). Due to the doubling of training time, we deemed this increase unnecessary. The following metrics were used to determine model performance: AUC, F1 score, accuracy, and precision. The best classification threshold was determined by the finding at which F1 score was highest (the point at which accuracy and precision intersect). Scores were averaged over 3 random folds.

**Fig. 3.**
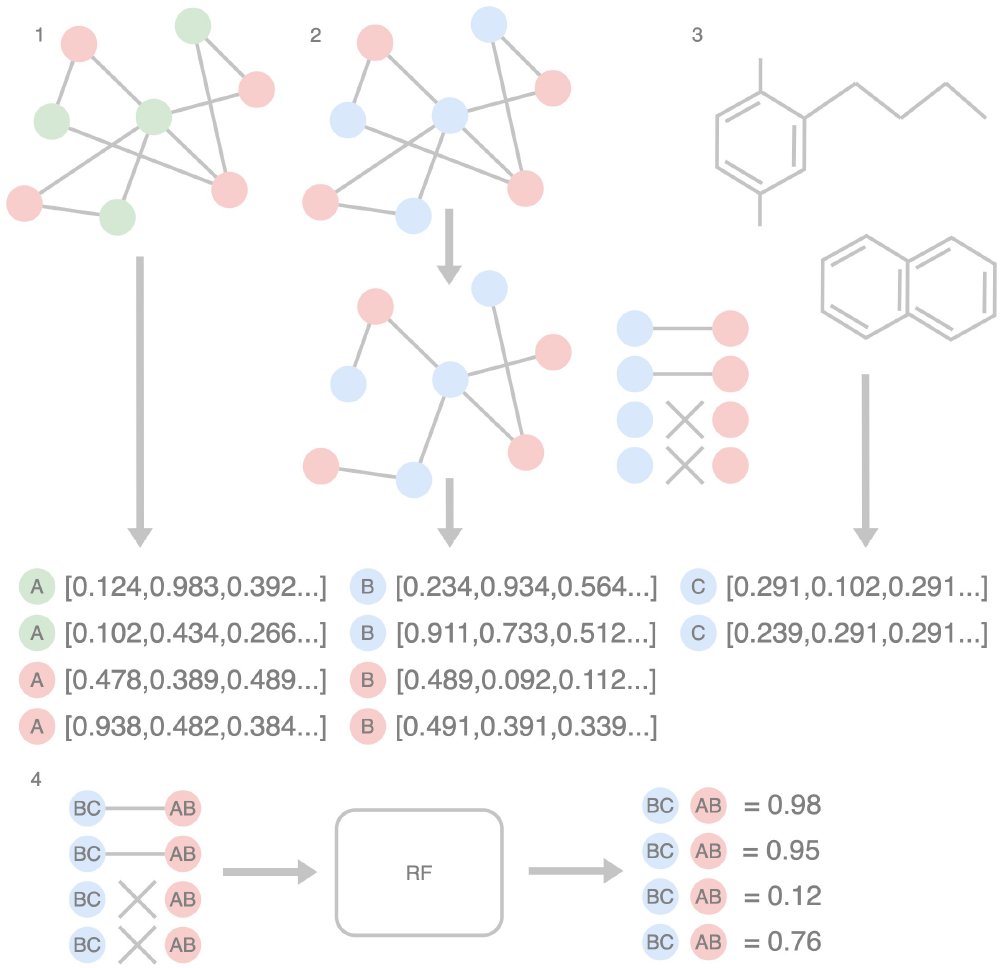
Workflow for model generation for *drug-inhibits-disease* edge. First, we split the graph into two subgraphs, a disease-gene subgraph (Red and green nodes, respectively), and a disease-mol subgraph (red and blue, respectively). 1) We generated embeddings for disease and gene nodes (Embeddings A). 2) We removed a subset of the edges of the disease-mol subgraph and generate node embeddings for disease and molecules (Embeddings B). 3) We generated embeddings based on the molecular substructure (Embeddings C). We now have two embeddings for each disease, and two for each molecule. To summarise, Embeddings A describe the neighbourhood or proximity of disease and genes in a disease-gene regulatory network (how close is one disease to another). Embeddings B describes the neighbourhood or proximity of a disease and molecule in a disease-mol regulatory network. Embeddings C describe the physicochemical similarity of compounds. 4) We combined the two embeddings for each disease and molecule (AB and BC, respectively), and used the edges removed from the disease-mol network to train a model capable of predicting the existence of a link between a disease and a molecule. In the example below, we can see there were two known disease-gene pairs (the regulation has been stated in Pathway Studio), and two random disease-gene pairs. We can see that the model has predicted the final pair to actually be an unreported regulation link.

For the protein-drug regulation prediction, node neighbourhood embeddings of dimension *d* for source and target *n*_*s,t*_, were complemented with structural information: *e*_*s*_ protein ProtVec amino acid trimer embeddings, and *e*_*t*_ Mol2Vec Morgan fingerprint substructure embeddings, where:

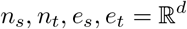

Each pair of embeddings was combined via one of various functions to create an edge embedding, before the edge embeddings of source and target were combined:

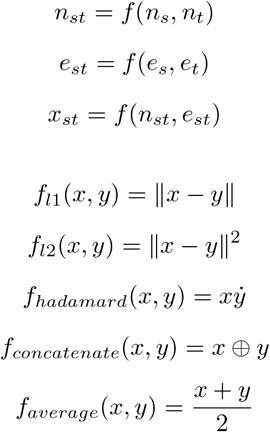

## Acknowledgments

The authors would like to thank Dr Ted Slater, Dr Mark Haupt and Dr Bruce Aronow. Without their contributions and insights, this manuscript would not be possible. Some contributors were afforded the time to perform this analysis by Elsevier, whom we also express our gratitude. Analysis was performed on the Elsevier Entellect machine learning platform, whose team we thank for their computational resource. The platform ingested the tremendously useful Pathway Studio, to which we thank Dr Anton Yuryev. Methodology was developed with the expert guidance of Dr Pan Pantziarka and Dr Javad Nazarian. This work is a follow up piece to the drug repurposing datathon between Mission:Cure, Elsevier and Pistoia Alliance. None of this would have been possible without the work of Dr Vladimir Makarov, to whom we express our greatest gratitude.

## Notes

### Competing Interest Statement

The authors have declared no competing interest.

